# Immunomodulatory effect of HLA-G Overexpressed Mesenchymal Stromal Cell in Cell-based Therapy for Myocardial Infarction

**DOI:** 10.1101/2025.07.02.662882

**Authors:** Wei Zhu, Jie Kong, Hong-Xia Li, Ting-Bo Jiang, Si-Jia Sun, Cao Zou

**Author notes:** Address for Correspondence: Cao Zou, MD, PhD or Si-Jia Sun, MD, PhD, Division of Cardiology, Department of Medicine, The First Affiliated Hospital of Soochow University, No. 188 Shizi Street, Suzhou, China. These authors have contributed equally to this work (Wei Zhu, Jie Kong).

## Abstract

**Background:** Our previous study demonstrated that intravenous administration of mesenchymal stromal cells (MSCs) significantly increased local cell engraftment and improved heart function. We sought to investigate whether HLA-G1 overexpressed MSCs could further increase local transplanted cells engraftment and improve heart function.

**Methods and Results:** Mice were randomized to receive intravenous administration of saline, human umbilical cord blood derived MSCs (hUCB-MSCs) 7 days prior to acute myocardial infarction (AMI), induced by ligation of the left anterior descending coronary artery. Then, intramyocardial transplantation of human induced pluripotent stem cell derived cardiomyocytes (hiPSC-CMs) was performed 30 minutes following AMI. Echocardiographic assessment was performed to assess heart function. *In-vivo* fluorescent imaging analysis were used to analyze cell engraftment. Flow cytometry of splenic regulatory T cells (Tregs) and natural killer (NK) cells was conducted to evaluate the immunomodulatory effect. Our result showed that systemic intravenous administration of hUCB-MSCs significantly increased systemic Tregs, decreased systemic NK cells, increased cell engraftment of intramyocardial transplanted hiPSC-CMs, culminating in improvement of heart function. Our in-vitro study showed that HLA-G1 overexpressed hUCB-MSCs modulated immune response by decreasing pro-inflammatory cytokines.

**Conclusions:** Systemic intravenous administration of HLA-G1 overexpressed hUCB-MSCs modulated immune response and enhanced the survival of local transplanted hiPSC-CMs to improve heart function following AMI.

## Introduction

Myocardial infarction (MI), which is characterized by imbalanced coronary blood perfusion and myocardial oxygen demand, place a heavy medical, social and financial burden on global economies^1^. Although advances achieved in revascularization treatment have improved the survival of MI following the primary ischemic insult, the incidence of post-MI heart failure (HF) has increased^2^. The infarcted hearts undergo a process of remodeling including disproportionate thinning of the infarcted myocardium and hypertrophy of non-infarcted myocardium, culminating in post-MI HF^3^. Current clinical pharmacological therapies as well as clinical application of cardiac assist devices and cardiac resynchronization therapy have limited the deterioration in cardiac function, reversed progression to post-MI HF and reduced mortality^4^. Nevertheless, current pharmacological strategies cannot replenish the lost cardiomyocytes (CMs). Regenerative therapies, seeking to regenerate and repair ischemic heart by administering exogenous cells, have been investigate as a promising strategy in cardiovascular disease^5^. Direct remuscularization with exogenous cells or activation of endogenous repair process by paracrine activities are two main targets in current cell-based therapies^6^. Although MI patients could be benefit from cell-based therapies, immunological rejection is an evitable barrier that limits clinical translation of cell-based regenerative therapies^7^.

There are several immune tolerance strategies available, including induction of central and peripheral immune tolerance, immunosuppressive agents, bioengineered cell patches and generation of universal cells^8–12^. Recent studies showed that induction of peripheral immune tolerance was one of the promising approaches to improve cell retention^8^. Our previous study showed that intravenous administration of human mesenchymal stromal cells (MSCs) could improve the survival and therapeutic efficacy of intramyocardial and intramuscular transplanted cells in both MI model and hind-limb ischemia model^13–14^. Recent studies demonstrated that human leukocyte antigen (HLA)-G modulated immune system though inducing HLA-G^+^ regulatory T cells (Tregs) and Foxp3^+^ Tregs as well as regulating natural killer (NK) cells^15^. HLA-G was one of the key molecules regulating maternal-fetal tolerance. Therefore, in this study, we hypothesized that intravenous systemic administration of HLA-G overexpressed MSCs could increase the cell retention and improve the therapeutic efficacy of intramyocardial transplanted human induced pluripotent stem cell (hiPSC)-CMs in a mouse model of MI. The underlying mechanism of immunomodulatory effect of HLA-G overexpressed MSCs was also investigated.

## Methods

This study was approved by the Ethics Committee of the First Affiliated Hospital of Soochow University (Project number 2022-159).

### Culture and preparation of human umbilical cord blood derived mesenchymal stromal cells and human induced pluripotent stem cell derived cardiomyocytes

Human umbilical cord blood derived MSCs (hUCB-MSCs) were used in this study (RC-005, Shenzhen Cell Inspire Biotechnology, Guangdong, China). hUCB-MSCs were purified by the MoFlo Cell sorting system (DxFLEX, Beckman Coulter, FL, USA). Sorted cells were characterized by the expression of CD90, CD105 and CD73; and absence of CD14, CD19, CD34, CD45 and HLA-DR with the ability to differentiate into osteoblasts, adipocytes and chondroblasts. Passage 4-6 hUCB-MSCs were used in the following steps. 5×10^5^ hUCB-MSCs were prepared in 100 μL of normal saline and intravenous administrated though the tail vein.

Human iPSC-CMs were used in this study (B0388-iCMs, Shanghai Amplicon-gene Bioscience, Shanghai, China). Beating clusters of hiPSC-CMs were observed under microscope. Immunostaining of cTnT and α-actinin 2 were used to confirm the expression of cardiomyocyte biomarkers and flow cytometry analysis of human cTnT showed that cTnT positive hiPSC-CMs accounted for 95.15 percentage of total cells. Before intramyocardial transplantation, hiPSC-CMs were stained with 5 μg/mL DiR cell-labeling solution (KGE2603-5, KeyGEN BioTECH, Jiangsu, China) for 15 min at 37 °C, washed and re-suspended in 30 μl of fresh plain medium for intramyocardial transplantation.

### Human HLA-G1 transfection

In this study, the HLA-G1 overexpression lentiviral vectors and the negative control vectors were obtained from Shanghai Genechem Co., Ltd. (Shanghai, China). hUCB-MSCs were transfected with HLA-G1 overexpression lentiviral vectors or negative control vectors to generate HLA-G1 overexpressed hUCB-MSCs or control hUCB-MSCs. In brief, hUCB-MSCs were cultured in 6-well plates (1×10^4^ cells/ml) for one day. Then, 20μL lentivirus with or without HLA-G1 gene (1×10^8^ TU/mL) were added into cell cultural medium. To evaluate the transfection efficacy of HLA-G1 overexpressing lentivirus, hUCB-MSCs were divided into three groups: (1) MSC group: hUCB-MSCs without transfection; (2) MSC-NC group: hUCB-MSCs transfected with GFP labelled lentivirus; (3) MSC-HLA-G1 group: hUCB-MSCs transfected with GFP labelled HLA-G1 overexpressing lentivirus. Co-cultured hUCB-MSCs with lentivirus with or without HLA-G1 gene for 24h, washed hUCB-MSCs and cultured the cells with fresh cultural medium for next 48h. HLA-G1 overexpressed hUCB-MSCs were verified by qRT-PCR and western blot.

### Establishment of mouse MI model, grouping and cell tracing

In this study, ICR mice were employed as their pro-inflammatory response in pro-inflammatory environment which contributed to evaluate the anti-inflammatory effect of administrated MSCs^16^. Intravenous administration of 100μL saline or 5×10^5^ hUCB-MSCs or 5×10^5^ HLA-G1 overexpressed hUCB-MSCs via the tail vein was performed one week prior to acute myocardial infarction (AMI) induction. AMI induction was conducted by the left anterior descending coronary artery ligation as described in our previous study^13^. Briefly, all the mice were anesthetized by intramuscular administration of tiletamine and zolazepam (Zoletil 50 mg/kg). Then, animals were ventilated and thoracotomy were performed between the third and fourth costae. Next, the left anterior descending coronary artery ligation were performed using 8-0 sterile silk suture in MI induction groups. 30 min after AMI induction, animals were randomized to intramyocardial administration of 30μL culture medium or 1×10^6^ DiR-labeled hiPSC-CMs at three different sites at the peri-infarct region. Subcutaneous administration of 200000U penicillin was performed for 5 days following AMI induction to prevent infection.

Mice which were died during MI induction procedure were excluded from the analysis (n=5). In this experiment, 30 male ICR mice, aged 6 weeks and weight 18-22g were included in the final analysis and randomized into 5 groups. (1) Sham group (n=6): intravenous administration of 100 μ L saline and thoracotomy were performed in this group; (2) AMI group (n=6): intravenous administration of 100μL saline, AMI induction and intramyocardial injection of 30 μL cultural medium were performed in this group; (3) AMI+CM group (n=6): intravenous administration of 100μL saline, AMI induction and intramyocardial administration of 1×10^6^ DiR-labeled hiPSC-CMs were performed in this group; (4) MSC+AMI+CM group (n=6): intravenous administration of 5×10^5^ hUCB-MSCs, AMI induction and intramyocardial administration of 1×10^6^ DiR-labeled hiPSC-CMs were performed in this group; (5) HLA-G1 MSC+AMI+CM group (n=6): intravenous administration of 5×10^5^ HLA-G1 overexpressed hUCB-MSCs, AMI induction and intramyocardial administration of 1×10^6^ DiR-labeled hiPSC-CMs were performed in this group. On day 28, all the animals were sacrificed by over anesthetized with 120mg/kg Zoletil after cardiac function assessment by transthoracic echocardiogram system (FUJIFILM Visual Sonics, Shanghai, China). Cell retention analysis by an IVIS® Spectrum optical imaging system (PerkinElmer, Inc. MA, USA) on day 7 and day 28. In brief, all the animals were anesthetized for epi-fluorsecent imaging using an IVIS® Spectrum optical imaging system under field of view (FOV) 10×10 cm (PerkinElmer, Inc. MA, USA). The injected DiR labeled cells were visualized under 750 nm/800 nm excitation/emission wavelength; and fluorescent intensity represented by radiant efficiency.

### Echocardiographic examination

Serial echocardiographic examination was performed by transthoracic echocardiogram system (FUJIFILM Visual Sonics, Shanghai, China) to assess cardiac function on day 28 after AMI induction. Before echocardiographic assessment, mice were anesthetized with isoflurane inhalation. Then, all echocardiographic assessment were performed by an experienced operator according to the European Society of Cardiology recommendations^17^. Left ventricular end-systolic dimension (LVESD), left ventricular end-diastolic dimension (LVEDD), fractional shortening (FS) and left ventricular ejection fraction (LVEF) were recorded to assess cardiac function and the mean value of three cardiac cycles was calculated for each animal.

### Histological Assessment

After all the animals sacrificed, hearts were harvested for the following histological assessment. The harvested tissues were fixed in ice-cold 4% buffered formalin, followed by dehydrated in the gradient ethanol and embedded in paraffin, then sectioned into 5μm slices. H&E staining was performed using the commercial kit (KGE1204-50, KeyGEN BioTECH, Jiangsu, China). Trichrome Masson’s staining was performed to determine the infarct size of the heart using Masson’s staining kit (KGE1113-8, KeyGEN BioTECH, Jiangsu, China). Infarct size was represented by the ratio of infarct area (characterized by fibrotic tissue strained blue) to LV area (characterized by myocardial tissue stained red and fibrotic tissues stained blue) and measured using Automated Microscope System (IXplore IX85 Pro, OLYMPUS, Tokyo, Japan) at three cross sections. The mean value of the three sections was calculated to represent infarct size. Immunohistochemical staining was performed to evaluate apoptosis in the peri-infarct area using a TdT-mediated dUTP Nick-End Labeling (TUNEL) kit (KGE1401-20, KeyGEN BioTECH, Jiangsu, China). Immunofluorescence staining of α-smooth muscle antigen (SMA, 1:200, 53-9760-82, Invitrogen, ThermoFisher, Shanghai, China) and von Willebrand factor (vWF, 1:200, PT0316R, Immunoway, Suzhou, China) were performed to assess neovascularization. Vessel density was represented as number of positive vessels per mm^2^ in the peri-infarct area. All data were measured in six random 40X fields from three different cross sections in a blinded fashion. Images of all sections were captured and analyzed by Automated Microscope System (IXplore IX85 Pro, OLYMPUS, Tokyo, Japan).

### Flow Cytometry Analysis

After all the mice were sacrificed, the splenocytes were harvested for the following flow cytometry analysis. In brief, single cell suspension of splenocytes were prepared in phosphate buffered saline (PBS) with the red blood cells removed. Then, one million splenocytes were blocked with the purified anti-mouse CD16/32 antibody (1:100, 101302, BioLegend, CA, USA) and 7AAD (1:50, 559925, BD Bioscience, NJ, USA) in flow buffer containing PBS, 1% BSA, 2 mM EDTA, 0.01% NaN3. Flow cytometry analysis of T cells and NK cells was performed as our previous study^13^. In brief, anti-mouse CD3 (1:100, 14-0032-82, Invitrogen, CA, USA), anti-mouse CD4 (1:100, 47-0042-82, Invitrogen, CA, USA) and anti-mouse CD8 (1:100, 11-0081-82, Invitrogen, CA, USA) was used for staining of CD3^+^, CD4^+^ and CD8^+^ T cells, respectively. Anti-mouse CD49b (1:100, 11-5971-82, Invitrogen, CA, USA) and anti-mouse NK1.1 (1:100, 17-5941-82, Invitrogen, CA, USA) were used for NK cell staining. Mouse Treg kit (88-8118-40, Invitrogen, CA, USA) including anti-mouse CD4 (1:100), anti-mouse CD25 (1:20) and anti-mouse Foxp3 (1:20) were used for Treg staining. Stained splenocytes were analyzed by a flow cytometer (CytoFLEX, Beckman Coulter, FL, USA) and fluorescence minus one (FMO) control was used for multicolor flow cytometry gating strategy. FlowJo software (V10.7.1, Becton, USA) was used for flow images analysis.

### The changes of cytokines

*In-vitro* co-culturing of CD4+ splenocytes with hUCB-MSCs or HLA-G1 overexpressed hUCB-MSCs were performed to assess the immunomodulatory effect of hUCB-MSCs. CD4^+^ cells were isolated by labeling the cells with anti CD4-FITC antibody for 30 min on ice and subsequently selected by fluorescence-activated cell sorting (FACS, CytoFLEX, Beckman Coulter, FL, USA). Then, 5×10^5^ CD4+ splenocytes were co-cultured with 1×10^5^ hUCB-MSCs or HLA-G1 overexpressed hUCB-MSCs for 7 days in transwell permeable supports (multiple well plate, Corning Life Science, MA, USA). Supernatant was collected for cytokine analysis using multiplex cytokine quantitation array as per the manufacturer’s instructions (Ray Biotech, Norcross, GA, USA).

### Statistical analysis

All data were analyzed by SPSS Statistics (version 27.0, IBM Corporation, Armonk, NY, USA). The results are presented as the mean±standard error of the mean (SEM) (continuous variables and normally distribution). For multiple groups analysis, one-way ANOVA with Tukey post hoc test were performed in a blind manner. A two-sided p<0.05 was regarded as a statistically significant difference.

## Results

In this study, five mice were died after AMI induction and in the final experiment, thirty mice were randomized into one of five groups: the Sham group (n=6), the AMI group (n=6), the AMI+CM group (n=6), the MSC+AMI+CM group (n=6) and the HLA-G1 MSC+AMI+CM group (n=6) (**Figure 1**). hUCB-MSCs were obtained from the Shenzhen Cell Inspire Biotechnology and flow cytometry analysis was performed to evaluate biomarker expression of hUCB-MSCs. Our results showed that hUCB-MSCs were expressing CD90, CD105 and CD73 as well as absence of CD14, CD19, CD34, CD45 and HLA-DR (**Supplemental Table 1**). Our results demonstrated that both GFP labelled lentivirus and GFP labelled HLA-G1 overexpressing lentivirus could successfully transfect hUCB-MSCs and after transfected with GFP labelled HLA-G1 overexpressing lentivirus, the protein level of HLA-G1 was significantly increased in the MSC-HLA-G1 group, compared with the MSC and the MSC-NC groups (**Supplemental Figure 1**). Immunofluorescence staining of cTnT and α-actinin was performed to evaluate the biomarker expression of hiPSC-CMs (**Supplemental Figure 2A&B**). Our flow cytometry result showed that 95.19% hiPSC-CMs were expressing cTnT (**Supplemental Figure 2C**). No tumor formation was observed at any injection site or organs. An elevated ST segment in the mouse electrocardiographic assessment was observed to confirm success AMI model establishment during AMI induction (**Supplemental Figure 3**).

**Figure 1.**
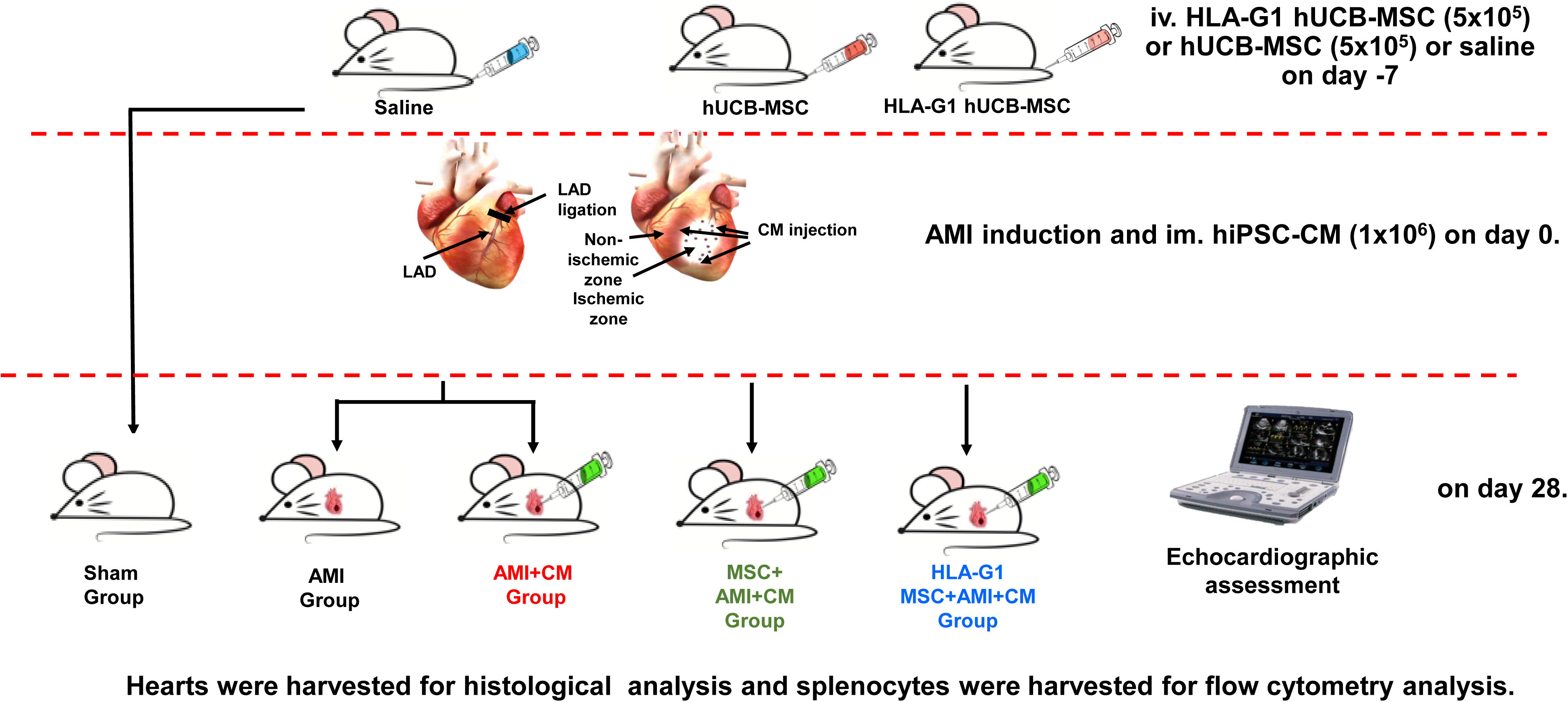
Flow chart of the experiment.

### Improved LV systolic function after transplantation

Standard transthoracic echocardiographic assessment was performed at the end of experiment and LVEF, FS, LVEDD, LVESD were recorded to evaluate cardiac function (**Figure 1A**).

Compared with the Sham group, LVEF (82.29±0.64 vs. 21.62±8.14) and FS (vs.) significantly decreased (**Figure 1B&1C**), whereas LVESD and LVEDD significantly increased in the AMI group (**Figure 1D&1E**). After hiPSC-CM transplantation, LVEF and FS were remarkedly improved (**Figure 1B&1C,** all p<0.05) and LVESD and LVEDD were remarkedly reduced in the AMI+CM, MSC+AMI+CM or HLA-G1 MSC+AMI+CM groups (**Figure 1D&1E,** all p<0.05). Consistently with our previous study, intravenous administration of MSCs could notably improve cardiac function in the MSC+AMI+CM or HLA-G1 MSC+AMI+CM groups (**Figure 1B, 1C, 1D&1E,** all p<0.05). Interestingly, intravenous administration of HLA-G1 overexpressed MSCs could further improve LVEF and FS and decrease LVESD as well as LVEDD in the HLA-G1 MSC+AMI+CM group compared to the MSC+AMI+CM group (**Figure 1B, 1C, 1D&1E,** all p<0.05).

Taken together, our results showed that consistently with our previous study, intramyocardial transplantation of hiPSC-CM with or without intravenous administration of hUCB-MSC could significantly improve cardiac function in AMI animals. Moreover, intravenous administration of HLA-G1 overexpressed hUCB-MSCs further improved cardiac function compared to intravenous administration of hUCB-MSCs.

### Increased engraftment and cell retention after transplantation

Fluorescent imaging of hiPSC-CM transplanted animals was performed at day 7 and day 28 to evaluate cell engraftment and cell retention of DiR-labeled hiPSC-CM in the peri-infarct area after transplantation (**Figure 2A**). At day 7, compared with the AMI+CM group, intravenous systemic administration of hUCB-MSCs or HLA-G1 overexpressed hUCB-MSCs could significantly increase engraftment and retention of local transplanted hiPSC-CMs (**Figure 2B**, all p<0.05). Furthermore, intravenously administered HLA-G1 overexpressed hUCB-MSCs could further improve survival of local injected hiPSC-CMs (**Figure 2B**, all p<0.05). At day 28, the similar results were observed that systemic administration of hUCB-MSCs or HLA-G1 overexpressed hUCB-MSCs increased engraftment of DiR-labeled hiPSC-CMs and an increased cell retention was observed in the HLA-G1 MSC+AMI+CM group compared to the MSC+AMI+CM group (**Figure 2C**, all p<0.05).

**Figure 2.**
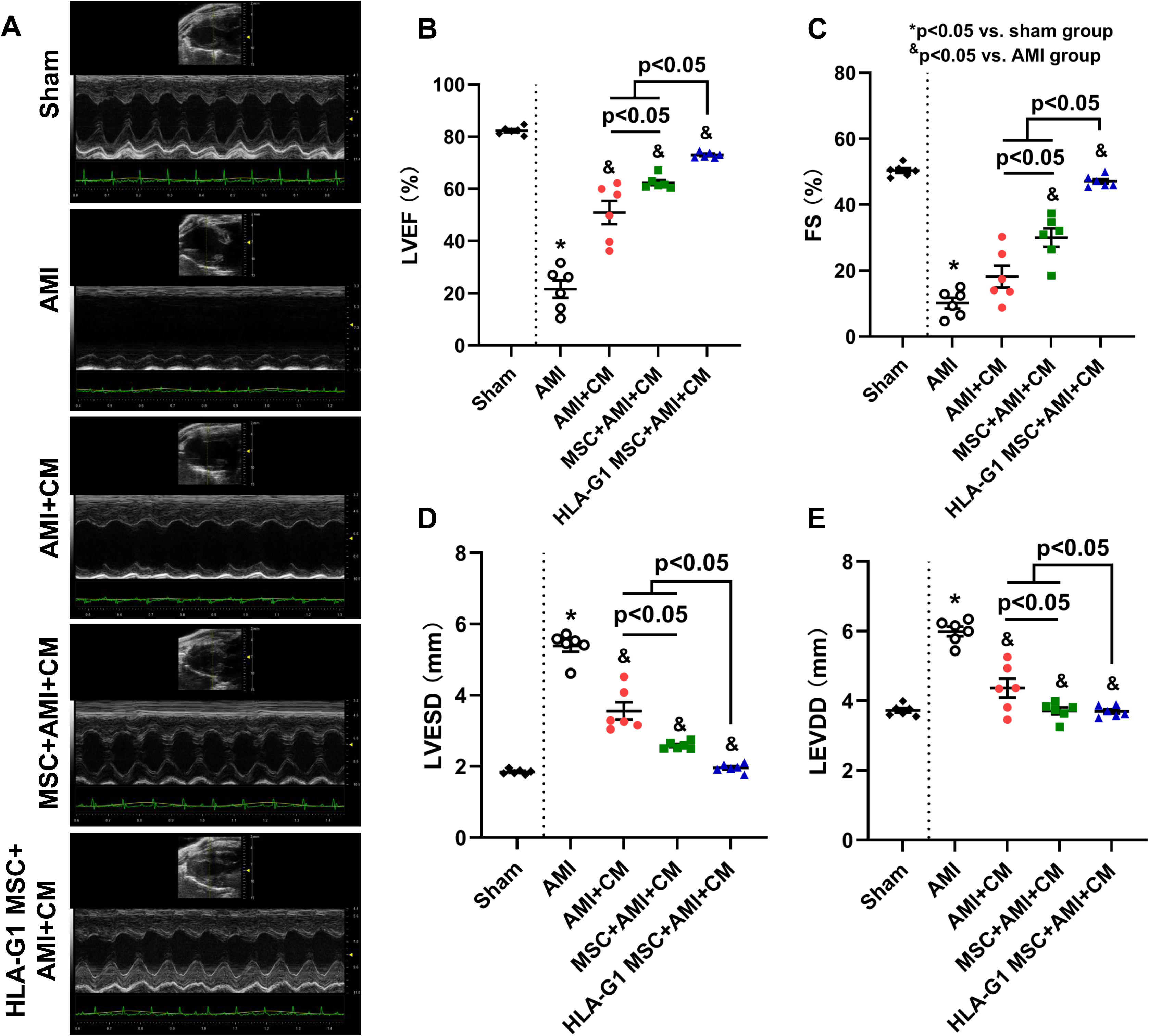
Intravenous systemic administration of HLA-G1 overexpressed hUCB-MSCs improved heart function. To assess heart function, echocardiographic assessment was performed (A) and left ventricular ejection fraction (LVEF) (B), fractional shortening (FS) (C), LV end-systolic dimension (LVESD) (D) as well as LV end-diastolic dimension (LVEDD) (E) were calculated. Intravenous administration of hUCB-MSCs or HLA-G1 overexpressed hUCB-MSCs could increase LVEF and FS as well as decrease LVESD and LVEDD. In addition, intravenous administration of HLA-G1 overexpressed hUCB-MSCs could further improve heart function compared with the MSC+AMI+CM group.

Taken together, our results demonstrated that consistent with our previous study, intravenous administration of hUCB-MSCs or HLA-G1 overexpressed hUCB-MSCs significantly improved the cell retention of intramyocardial transplanted hiPSC-CMs. The increased immunomodulatory effect of HLA-G1 overexpressed hUCB-MSCs was observed in the HLA-G1 MSC+AMI+CM group compared to the MSC+AMI+CM group.

### Decreased infarct size and apoptosis rate after transplantation

Masson trichrome staining was performed to evaluate the infarct size and immunohistological staining of TUNEL was performed to assess the apoptosis rate. The image of Masson trichrome staining, H&E staining and immunohistology staining of TUNEL were showed in **Figure 3A**. Significant infarct area and increased apoptosis were observed in the AMI group (**Figure 3B&3C**). Compared with the AMI group, intramyocardial transplantation of hiPSC-CMs with or without intravenous administration of hUCB-MSCs significantly reduced infarcted size and decreased apoptosis of ischemic cardiomyocytes in the peri-infarcted area (**Figure 3B&3C**). Consistent with our previous studies, intravenous administration of hUCB-MSCs improved the therapeutic efficacy of intramyocardial transplanted hiPSC-CMs (**Figure 3B&3C**). Moreover, intravenous administration of HLA-G1 overexpressed hUCB-MSCs further decreased infarcted size and apoptosis of ischemic cardiomyocytes in the peri-infarcted area (**Figure 3B&3C**).

**Figure 3.**
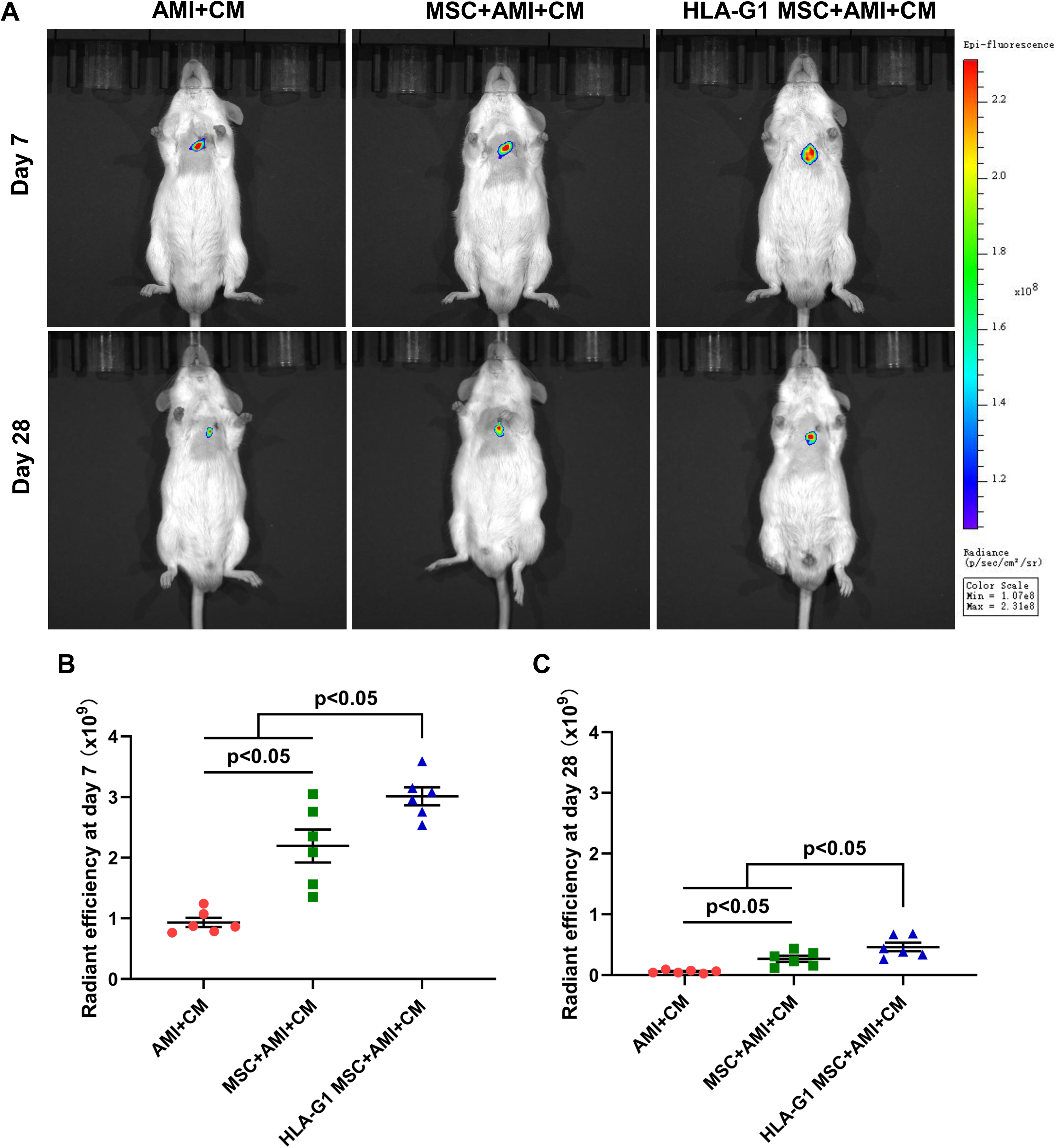
Intravenous administration of HLA-G1 overexpressed hUCB-MSCs could increase cell engraftment. To evaluate the cell engraftment of intramyocardial transplanted hiPSC-CMs, fluorescent imaging of DiR-labeled hiPSC-CM transplanted animals was performed at day 7 and day 28 (A). Compared with the AMI+CM group, intravenous administration of hUCB-MSCs significantly increased cell engraftment of intramyocardial transplanted hiPSC-CMs in the MSC+AMI+CM group on day 7 and 28 (B&C). Moreover, intravenous administration of HLA-G1 overexpressed hUCB-MSCs could further increase cell survival of transplanted hiPSC-MSCs in the HLA-G1 MSC+AMI+CM group compared with the MSC+AMI+CM group on day 7 and 28 (B&C).

Taken together, consistent with our previous study, intravenous administration of hUCB-MSCs could significantly reduced the infarct size and apoptosis of intramyocardial transplanted hiPSC-CMs. Moreover, overexpressing HLA-G1 in hiPSC-MSCs could improve the immunomodulatory effect of hiPSC-MSCs.

### Increased angiogenesis and neovascularization

Immunofluorescence staining of α-SMA and vWF were performed to evaluate the angiogenesis and neovascularization in the peri-infarct area (**Figure 4A**). Both α-SMA and vWF positive vessels significantly decreased in the AMI group compared to the Sham group and intramyocardial transplantation of hiPSC-CMs remarkably improved angiogenesis in the hiPSC-CM transplantation groups (**Figure 4B&C**). Intravenous administration of hUCB-MSCs increased angiogenesis in the peri-infarct area in the MSC+AMI+CM group compared to the AMI+CM group (**Figure 4B&C**). In addition, intravenous administration of HLA-G1 overexpressed hUCB-MSCs instead of hUCB-MSCs could further increased angiogenesis in the HLA-G1 MSC+AMI+CM group (**Figure 4B&C**).

**Figure 4.**
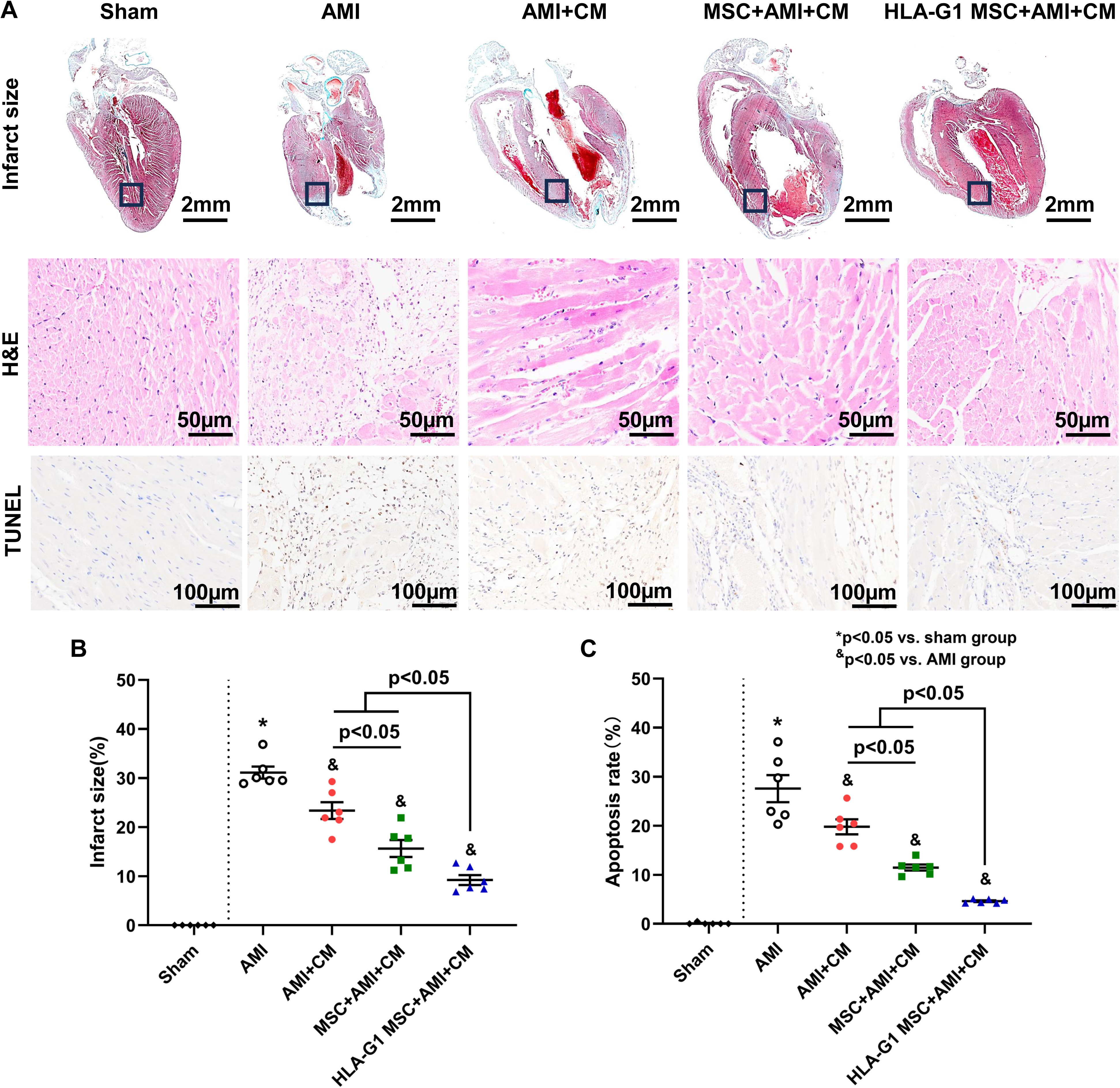
Intravenous administration of HLA-G1 overexpressed hUCB-MSCs decreased infarct size and apoptosis. Masson’s trichrome staining, H&E staining and immunohistology staining of TUNEL were performed to evaluate infarct size and apoptosis (A). Intravenous administration of hUCB-MSCs or HLA-G1 overexpressed hUCB-MSCs significantly decrease infarct size and apoptosis compared with the AMI and the AMI+CM groups (B&C). Moreover, intravenous administration of HLA-G1 overexpressed hUCB-MSCs in the HLA-G1 MSC+AMI+CM group further reduced infarct size and apoptosis, compared with the MSC+AMI+CM group (B&C).

Taken together, intravenous administration of hUCB-MSCs significantly improved the therapeutic efficacy of local transplanted hiPSC-CMs and angiogenesis in the peri-infarct area. In addition, an increased angiogenesis and neovascularization were observed in the HLA-G1 MSC+AMI+CM group.

### The immunomodulatory effect of hUCB-MSCs and HLA-G1 overexpressed hUCB-MSCs

To assess the immunomodulatory effect of hUCB-MSCs and HLA-G1 overexpressed hUCB-MSCs, fresh spleens were isolated and flow cytometry of splenocytes was performed (**Figure 5A**). There was no significant difference between the Sham, the AMI and the AMI-CM groups (**Figure 5B**). Nevertheless, intravenous administration of hUCB-MSCs or HLA-G1 overexpressed hUCB-MSCs significantly increased splenic Tregs in the MSC+AMI+CM and the HLA-G1 MSC+AMI+CM groups (**Figure 5B**). Moreover, intravenous administration of HLA-G1 overexpressed hUCB-MSCs further increased splenic Tregs in the HLA-G1 MSC+AMI+CM group compared to the MSC+AMI+CM group (**Figure 5B**). Splenic NK cells significantly increased after MI induction and intramyocardial transplanted hiPSC-CMs had no effect on splenic NK cells (**Figure 5C**). Intravenous administration of hUCB-MSCs or HLA-G1 overexpressed hUCB-MSCs significantly decreased splenic NK cells in the MSC+AMI+CM and the HLA-G1 MSC+AMI+CM groups (**Figure 5C**). Moreover, intravenous administration of HLA-G1 overexpressed hUCB-MSCs further reduced splenic NK cells in the HLA-G1 MSC+AMI+CM group compared to the MSC+AMI+CM group (**Figure 5C**). There was no significant difference between different groups on splenic CD4^+^ T cells (**Figure 5D**).

**Figure 5.**
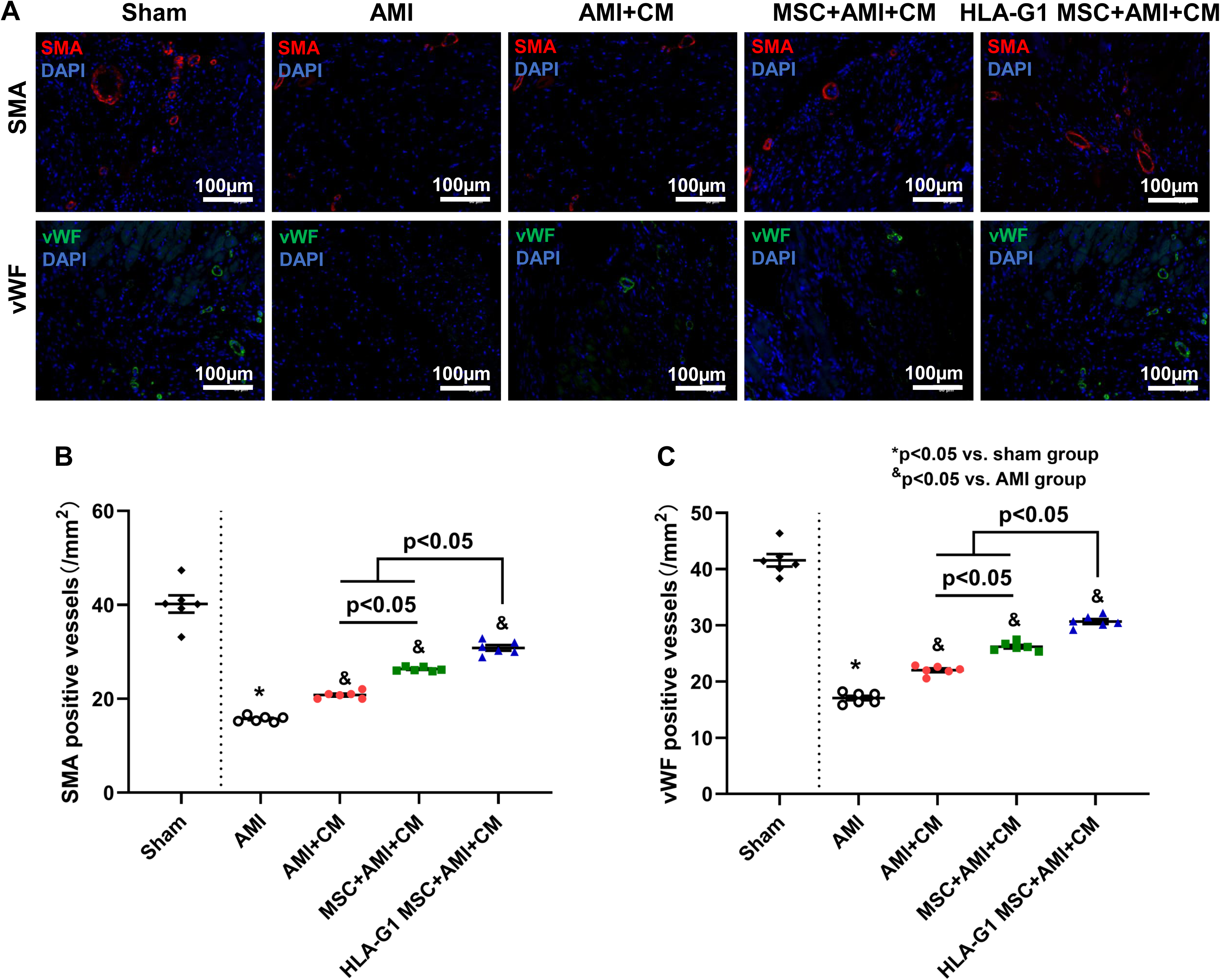
Intravenous administration of HLA-G1 overexpressed hUCB-MSCs increased neovascularization. Immunofluorescence staining of α-SMA and vWF were performed to assess neovascularization in the peri-infarct area (A). Compared with the AMI and AMI+CM groups, intravenous administration of hUCB-MSCs or HLA-G1 overexpressed hUCB-MSCs significantly increased angiogenesis (B&C). In addition, intravenous administration of HLA-G1 overexpressed hUCB-MSCs significantly increased angiogenesis in the HLA-G1 MSC+AMI+CM group, compared with the MSC+AMI+CM group (B&C).

In addition, the changes to cytokine profile were also measured 7 days after co-culturing of hUCB-MSCs or HLA-G1 overexpressed hUCB-MSCs with splenic CD4^+^ T cells. After co-cultured with hUCB-MSCs, the expression of pro-inflammatory cytokines including interferon-γ (IFN-γ), tumor necrosis factor-α (TNF-α), interleukin (IL)-2 and IL-17A significantly decreased, although anti-inflammatory cytokines remain unchanged (**Figure 6A-G**). Moreover, HLA-G1 overexpressed hUCB-MSCs further reduced pro-inflammatory cytokines compared to hUCB-MSCs (**Figure 6A-G**).

**Figure 6.**
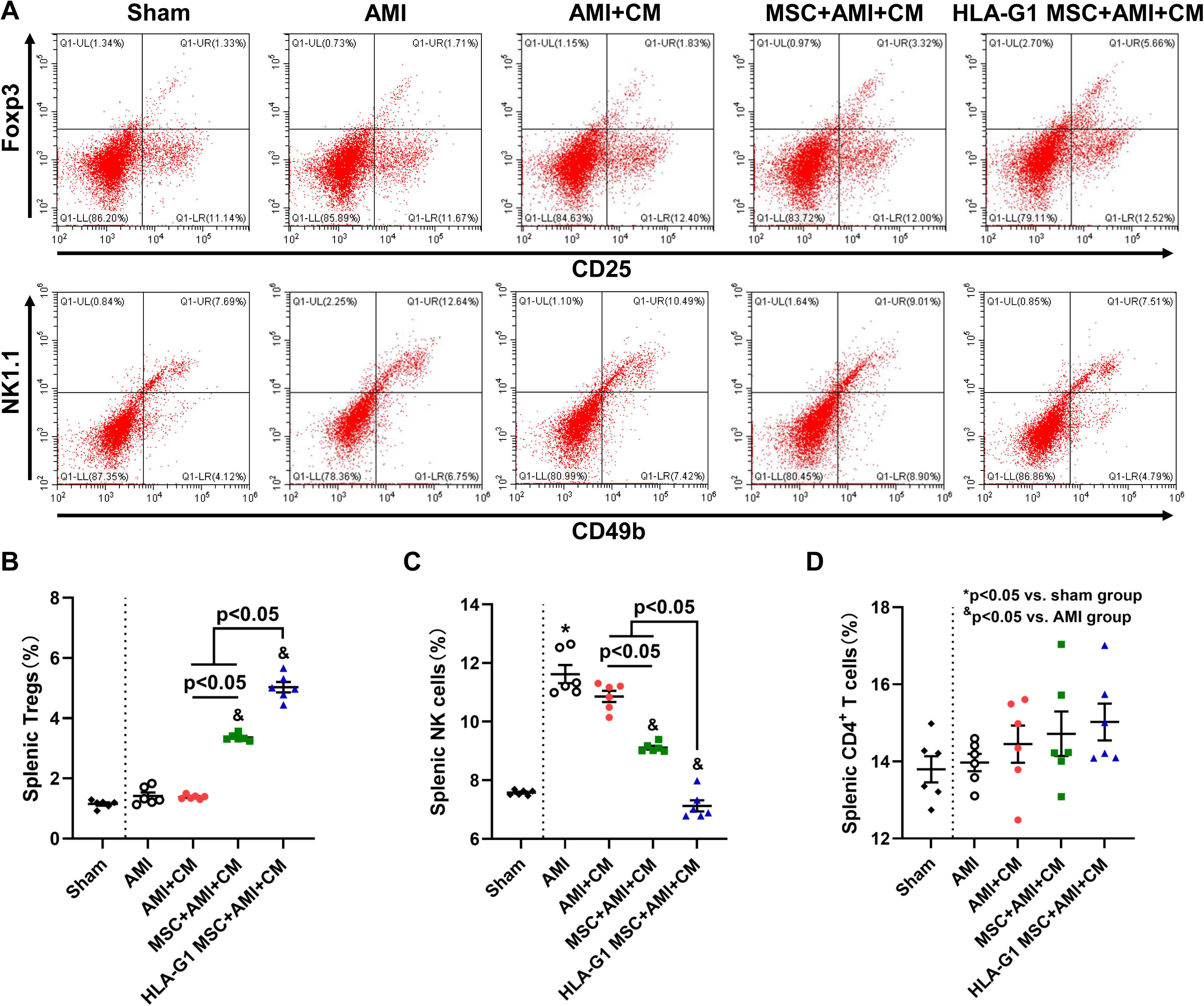
Intravenous administration of HLA-G1 overexpressed hUCB-MSCs increased Tregs and decreased NK cells. Flow cytometry analysis of splenocytes was performed to evaluate immunomodulatory effect of hUCB-MSCs (A). Intramyocardial transplantation of hiPSC-CMs has no effect on splenic Tregs and NK cells (B&C). Intravenous administration of hUCB-MSCs or HLA-G1 overexpressed hUCB-MSCs significantly increased splenic Tregs and decreased splenic NK cells (B&C). In addition, intravenous administration of HLA-G1 overexpressed hUCB-MSCs further increased splenic Tregs and decreased splenic NK cells compared with the MSC+AMI+CM group (B&C). Intravenous administration of hUCB-MSCs or HLA-G1 overexpressed hUCB-MSCs have no effect on splenic CD4^+^ T cells.

**Figure 7.**
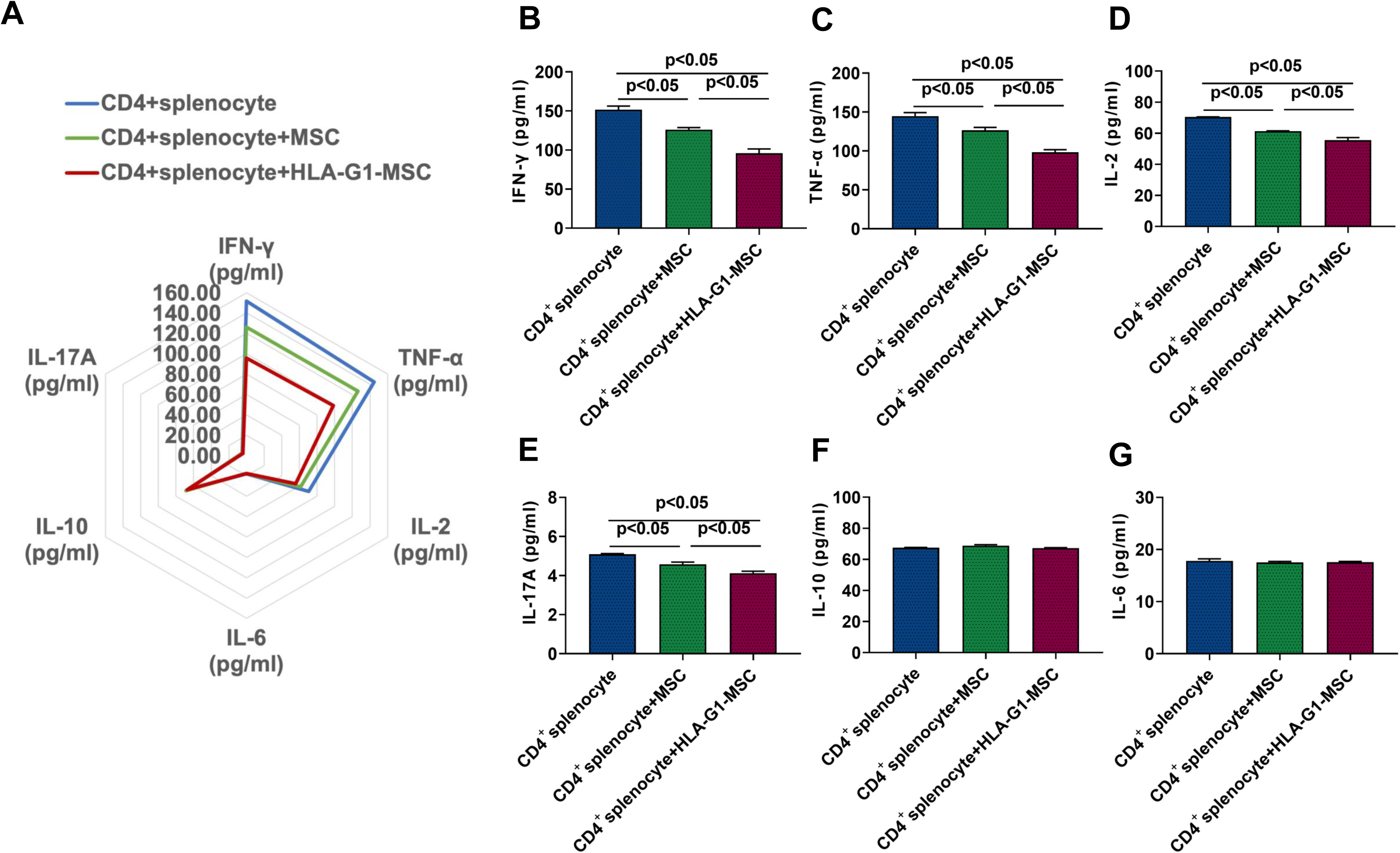
Cytokine profiles changes after co-culture for 7 days with CD4^+^ splenocytes and hUCB-MSCs or HLA-G1 overexpressed hUCB-MSCs. To assess the changes of cytokine profiles, the supernatant level of cytokines was measured at day 7 (A). interferon (IFN)-γ (B) and tumor necrosis factor (TNF)-α (C), interleukin (IL)-2 (D) and IL-17A (E) in the supernatant reduced significantly after co-culture with hUCB-MSCs for 7 days, compared with the cytokine level in the supernatant of the CD4^+^ cell population alone. HLA-G1 overexpressed hUCB-MSCs have superior immunomodulatory effects on these cytokines compared with hUCB-MSCs. The cytokine level of IL-10 (F) and IL-6 (G) remained unchanged.

Taken together, our results demonstrated that intravenous administration of HLA-G1 overexpressed hUCB-MSCs had superior immunomodulatory efficacy compared to intravenous administration of hUCB-MSCs. These immunomodulatory effects at least partily contributed to decreased pro-inflammatory cytokine release.

## Discussions

### Main findings

In this study, our results demonstrated that consisting with our previous study, intravenous administration of hUCB-MSCs significantly increased the survive of intramyocardial transplanted hiPSC-CMs. HLA-G1 overexpressed hUCB-MSCs have the superior immunomodulatory efficacy over hUCB-MSCs. The decreased pro-inflammatory cytokines and increased Tregs in the spleen contribute to the improved immunomodulatory effect and increased cell engraftment of intramyocardial transplanted hiPSC-CMs. In addition, the increased cell engraftment of intramyocardial transplanted hiPSC-CMs was associated further improved cardiac function, which is mediated with increased angiogenesis and decreased inflammation and apoptosis in the peri-infarct area.

### Immunomodulatory effects of HLA-G1 overexpressed hUCB-MSCs

A wide spectrum of studies have demonstrated that HLA-G is a non-classical HLA class I molecule widely expressed at the maternal-fetal interface and on several immuneprivileged tissues or organs and plays a vital role in the maternal-fetal immunotolerance^15, 18, 19^. Moreover, overexpression of HLA-G1 is widely used to reduce inflammation and increase immunomodulatory effect in cell transplantation^20^. Compared with other source derived MSCs, hUCB-MSCs expressed higher HLA-G1 molecules under standard cultivation^21^. Therefore, in this study, HLA-G1 overexpressed hUCB-MSC was employed to improve cell survival of intramyocardial transplanted iPSC-CMs. We intravenous administration of hUCB-MSCs or HLA-G1 overexpressed hUCB-MSCs 7 days prior to intramyocardial hiPSC-CM transplantation, as our previous study showed that splenic Tregs reached their peak level at 7 days following intravenous infusion of MSCs. Our results showed that intravenous administration of HLA-G1 overexpressed hUCB-MSCs further improved cell engraftment of intramyocardial transplanted hiPSC-CMs, compared with intravenous injection of hUCB-MSCs. These results supported that increased immunomodulatory effects were at least partly attributed to increased expression of HLA-G1. The increased expression of HLA-G1 was associated with increased splenic Tregs and decreased splenic NK cells. Previous studies showed that immunological recognition and rejection by the host were orchestrated by both innate and adaptive immune cells^22^. CD8^+^ cytotoxic T cells and CD4^+^ Th cells attack allogenic cells though major histocompatibility complex (MHC) class I molecules dependent manner, whereas NK cells were activated by MHC independent manner^22^. Our results showed that intravenous administration of HLA-G1 overexpressed hUCB-MSCs reduced immunological rejection though both MHC dependent and MHC independent manner, as both increased splenic Tregs and decreased NK cells were observed in our study. Indeed, improved cell engraftment was observed after intravenous systemic administration of HLA-G1 overexpressed hUCB-MSCs.

To further investigate the immunomodulatory effect of HLA-G1 overexpressed hUCB-MSCs, the changes of cytokines were analyzed after 7 days of co-culture with hUCB-MSCs and CD4^+^ splenocytes. Our previous study showed that intravenous administration of hiPSC-MSCs could significantly decrease IFN-γ that regulate immunological response, as well as pro-inflammatory cytokines including TNF-α and IL-17A^13^. Interesting, our *in-vitro* study demonstrated that apart from IFN-γ, TNF-αand IL-17A, intravenous administration of hUCB-MSCs or HLA-G1 overexpressed hUCB-MSCs could reduce pro-inflammatory cytokines IL-2. Previous studies showed that low-dose IL-2 promoted the proliferation and activation of regulatory cells, whereas high-dose IL-2 improved the immune response of effector cells^23^. In our study, decreased IL-2 did not lead to reduced Tregs. This is likely because MSCs secret multiply cytokines such as indoleamine 2, 3-dioxygenase (IDO) and HLA-G, which promote the activation and proliferation of Tregs^24^. Our previous study showed that intravenous administration of hiPSC-MSCs induced tolerance of isogenic iPSC-CMs derived from the same iPSC line^13^. In this study, our results supported that both hiPSC-MSC or hUCB-MSC could achieve the same immunomodulatory efficacy. Nevertheless, we did not perform comparison between hiPSC-MSC and hUCB-MSC. It is hard to determine which cell type have superior immunomodulatory effect over the other one.

Taken together, our study demonstrated that intravenous administration of hUCB-MSCs modulated immune response through decreasing pro-inflammatory cytokines and NK cells as well as increasing Tregs, culminating in improvement of intramyocardial transplanted hiPSC-CMs.

### Mechanism of cardiac function improvement

Our results showed that intravenous administration of hUCB-MSCs or HLA-G1 overexpressed hUCB-MSCs significantly improved cardiac function 28 days after MI and intravenous infusion of HLA-G1 overexpressed hUCB-MSCs had the superior therapeutic efficacy over hUCB-MSCs. These therapeutic effects were associated with decreased infarct size and apoptosis as well as increased cell engraftment and neovascularization. Previous studies demonstrated that both intravenous administration of hUCB-MSCs and intramyocardial transplantation of hiPSC-CM could benefit cardiac repairment and heart function^25–27^. Therefore, the therapeutic effects of cell-based therapy in this study were due to both systemic immunomodulatory effect of intravenous infused hUCB-MSCs and paracrine effect of intramyocardial transplanted hiPSC-CMs. No tumor was observed in any injected site or any other organs. No any transplantation related arrhythmia was observed 7 days or 28 days after hiPSC-CMs transplantation. The therapeutic effect of intramyocardial transplanted hiPSC-CM was prefer to their paracrine effect rather than direct myocardial regeneration. Our previous studies demonstrated that intramyocardial transplantation of hiPSC-MSCs or hiPSC-CMs with or without intravenous administration of hiPSC-MSCs could modulate cardiac repair and regeneration^13^. The results of current study further confirmed that intravenous administration of MSCs combined with intramyocardial transplantation of CMs improved cardiac function of infarct heart. In addition, overexpression of HLA-G1 in infused hUCB-MSCs further improved cardiac repair and function. As no tumor formation and no transplantation relevant arrhythmia were detected, our study supported that both intramyocardial transplantation of hiPSC-CMs and systemic intravenous administration of hiPSC-MSCs were safe for the treatment of cardiovascular disease. Taken together, the results of this study support the application of systemic intravenous administration of hUCB-MSCs and intramyocardial transplantation of hiPSC-MSCs to promote the repair and regeneration of infarct heart. Future studies could be launched to explore clinical application of systemic intravenous administration of hUCB-MSCs and intramyocardial transplantation of hiPSC-MSCs to repair the infarct heart and improve cardiac function.

### Limitation

First of all, our previous study showed that Tregs progressively increased after a single intravenous infusion of MSCs and reached their peak level 7 days after systemic intravenous administration of MSCs^13^. Therefore, we performed systemic intravenous infusion of MSCs 7 days prior to MI induction. Nonetheless, it is hard to mimic clinical scenario of MI patients. The optimal time of intravenous MSC infusion in clinical application desired to be further investigated. Secondly, our previous study demonstrated that multiple intravenous administration of MSCs achieved the superior therapeutic effect compared to a single intravenous injection^14^. Whether intravenous administration of hUCB-MSCs every 7 days could further improve the immunomodulatory effect in MI animals remain unclear. Nonetheless, our resulted showed that overexpression of HLA-G1 was an effective approach to improve the immunomodulatory effect of hUCB-MSCs. Thirdly, previous studies demonstrated that HLA-G molecules modulated NK cells though KIR2DL4-mediated signaling and increased Tregs though increasing the expression of Foxp3 in T cells^15, 28^. Our results confirmed the immunomodulatory effect of HLA-G molecules on NK cells and Tregs. Nevertheless, future studies are desired to be launched to investigate whether HLA-G1 overexpressed hUCB-MSCs modulate the immune system through the same pathways. Fourthly, our *in-vivo* study demonstrated that a significant increase of human HLA-G1 gene expression and human HLA-G1 protein expression in HLA-G1 overexpressed hUCB-MSCs. It is difficult to determine whether intravenous administration of HLA-G1 overexpressed hUCB-MSCs could increase systemic HLA-G molecule expression in mouse MI model, as there was lack of a clear HLA-G ortholog in mouse. Nonetheless, a significant decrease of pro-inflammation cytokines was observed in mouse splenic CD4^+^ cells 7 days after co-cultured with hUCB-MSCs. Our study provided solid evidence on the immunomodulatory effects of HLA-G1 overexpressed hUCB-MSCs.

## Conclusion

This study provide solid evidence on improved immunomodulatory effects on hUCB-MSCs with overexpressed HLA-G1 gene and intravenous administration of HLA-G1 overexpressed hUCB-MSCs increased cell engraftment of intramyocardial transplanted hiPSC-CMs as well as heart function though increasing systemic Tregs, decreasing systemic NK cells and modulating immune cytokines releasing.

## Nonstandard Abbreviations and Acronyms

AMI: Acute Myocardial Infarction
CM: Cardiomyocyte
FMO: Fluorescence Minus One
FS: Fractional Shortening
FOV: Field Of View
HF: Heart Failure
HLA: Human Leukocyte Antigen
hiPSC: Human Induced Pluripotent Stem Cell
hUCB: Human Umbilical Cord Blood
IDO: Indoleamine 2, 3-Dioxygenase
IFN: Interferon
IL: Interleukin
LVEDD: left ventricular end-diastolic dimension
LVEF: left ventricular ejection fraction
VESD: left ventricular end-systolic dimension
MHC: Major Histocompatibility Complex
MSC: Mesenchymal Stromal Cell
MI: Myocardial Infarction
NK: Natural Killer
PBS: Phosphate Buffered Saline
SEM: Standard Error of the Mean
SMA TNF: Smooth Muscle Antigen Tumor Necrosis Factor
Treg: Regulatory T cell
TUNEL: TdT-mediated dUTP Nick-End Labeling
vWF: Von Willebrand Factor<colcnt=2>

---

**Supplemental Figure 1.**
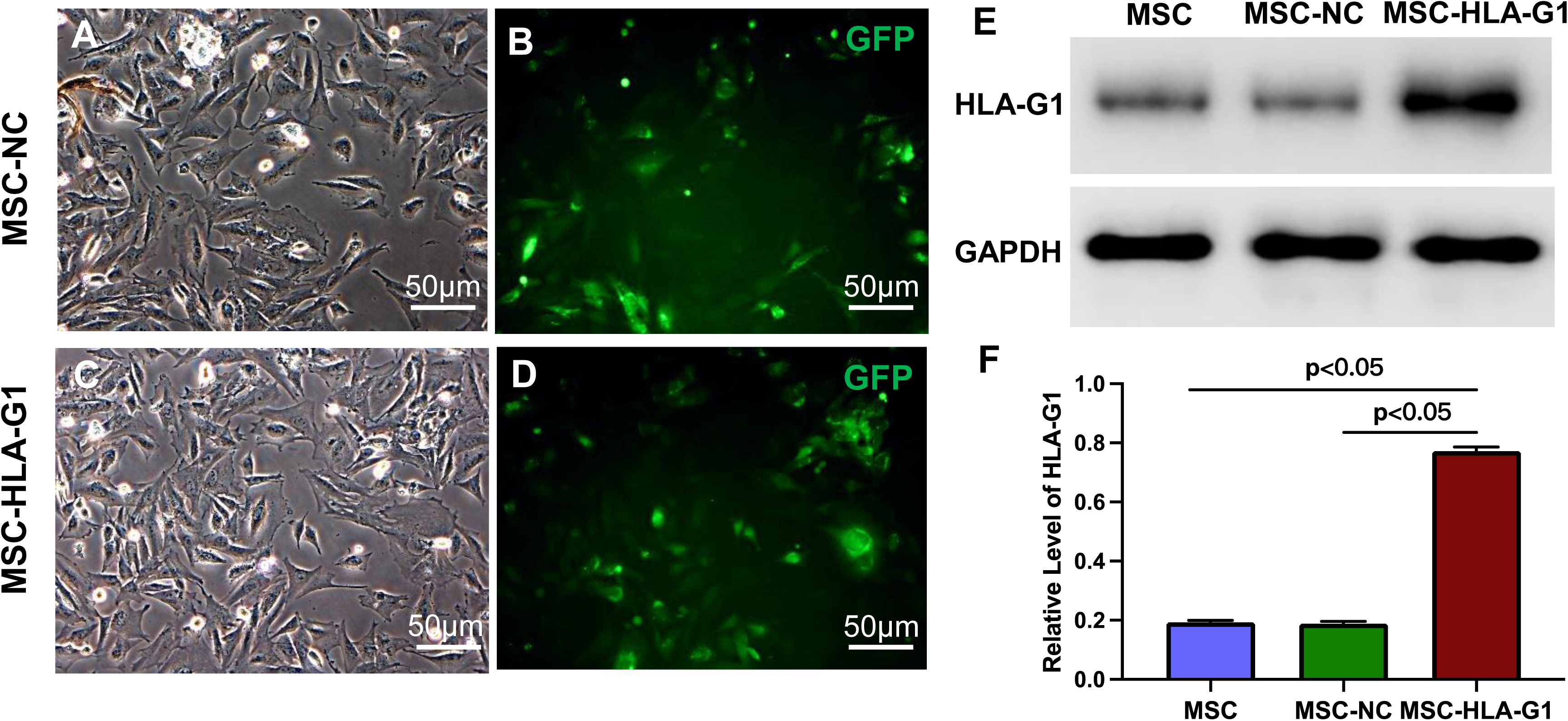
Establishment of HLA-G1 overexpressed hUCB-MSCs. Both GFP labelled lentivirus and GFP labelled HLA-G1 overexpressing lentivirus could successfully transfect hUCB-MSCs (A-D). Western blot was performed to detect protein expression of HLA-G1 (E). After transfected with GFP labelled HLA-G1 overexpressing lentivirus, the protein level of HLA-G1 were significantly increased in the MSC-HLA-G1 group, compared with the MSC and the MSC-NC groups (F).

**Supplemental Figure 2.**
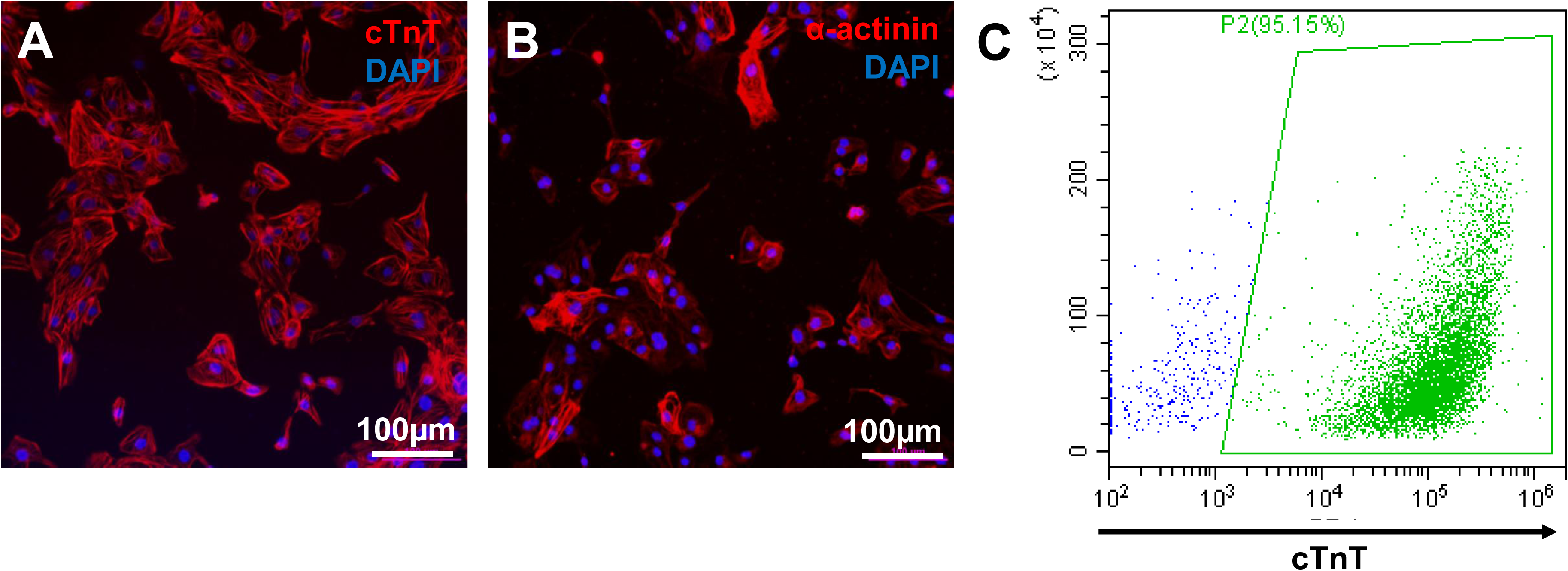
Biomarkers of hiPSC-CM. To detect biomarker expression of hiPSC-CMs, immunofluorescence staining of cTnT and α-actinin was performed (A&B). Our flow cytometry result showed that 95.19% hiPSC-CMs were expressing cTnT (C).

**Supplemental Figure 3.**
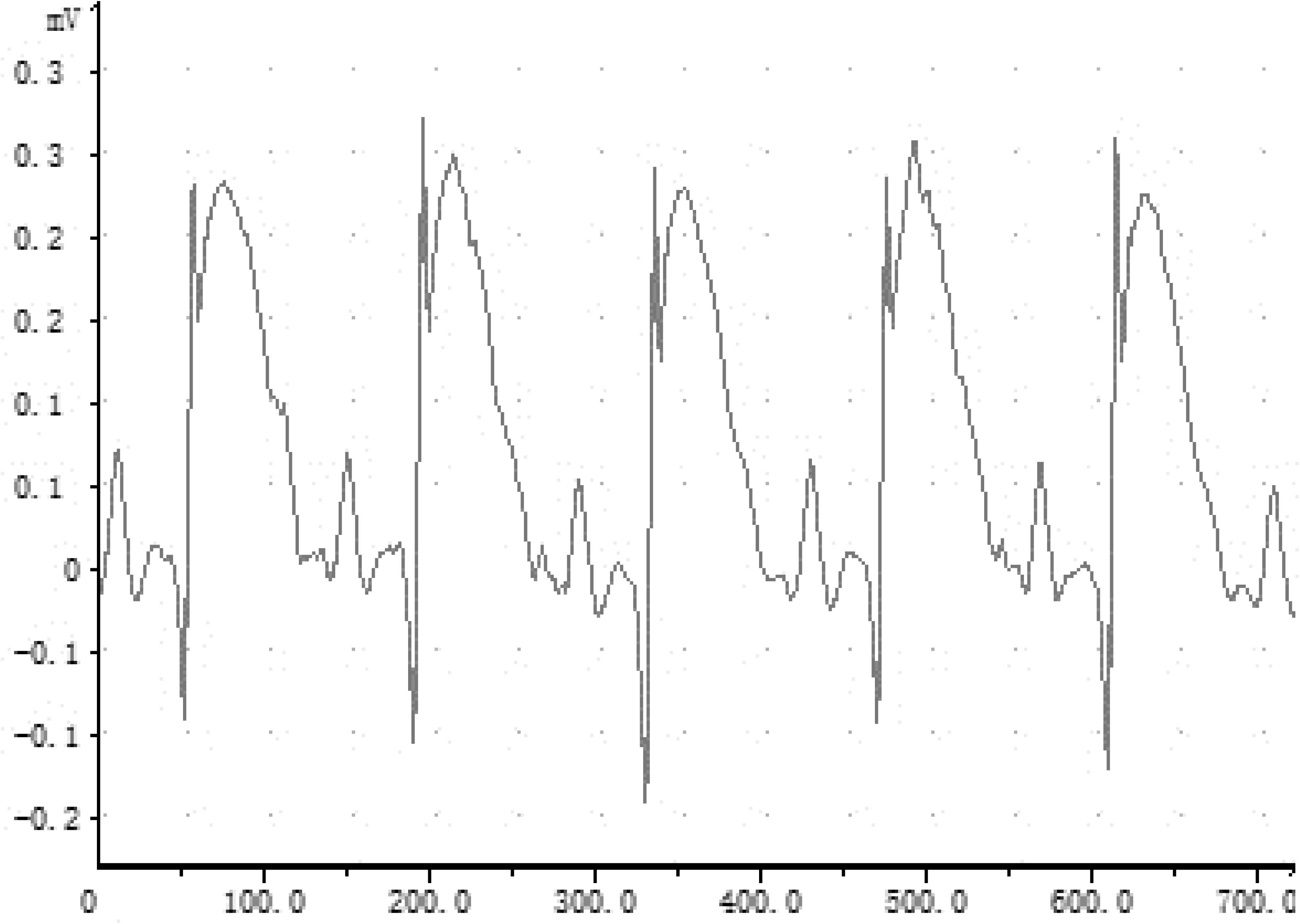
Electrocardiographic assessment after MI induction.

**Supplemental Table 1.**
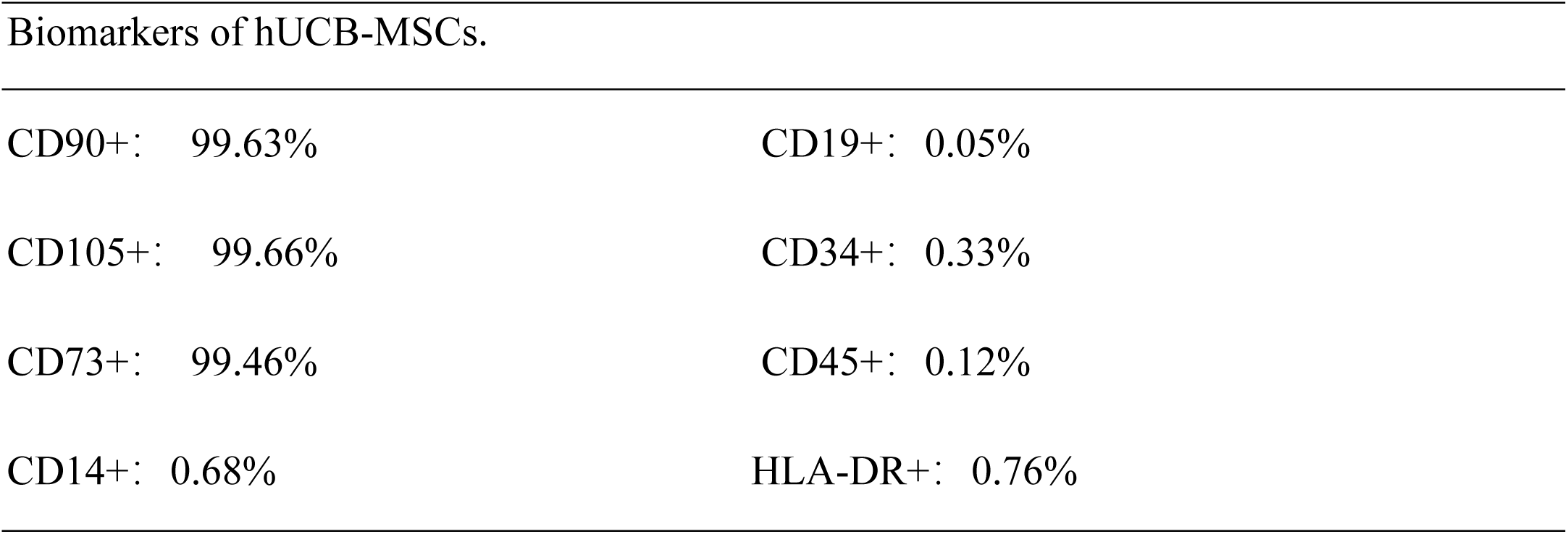
Biomarkers expression of hUCB-MSCs.

| Biomarkers of hUCB-MSCs. |  |
| --- | --- |
| CD90+: 99.63% | CD19+: 0.05% |
| CD105+: 99.66% | CD34+: 0.33% |
| CD73+: 99.46% | CD45+: 0.12% |
| CD14+: 0.68% | HLA-DR+: 0.76% |

## References

1. Martin SS, Aday AW, Allen NB, Almarzooq ZI, Anderson CAM, Arora P, Avery CL, Baker-Smith CM, Bansal N, Beaton AZ, Commodore-Mensah Y, Currie ME, Elkind MSV, Fan W, Generoso G, Gibbs BB, Heard DG, Hiremath S, Johansen MC, Kazi DS, Ko D, Leppert MH, Magnani JW, Michos ED, Mussolino ME, Parikh NI, Perman SM, Rezk-Hanna M, Roth GA, Shah NS, Springer MV, St-Onge MP, Thacker EL, Urbut SM, Van Spall HGC, Voeks JH, Whelton SP, Wong ND, Wong SS, Yaffe K, Palaniappan LP; American Heart Association Council on Epidemiology and Prevention Statistics Committee and Stroke Statistics Committee. 2025 Heart Disease and Stroke Statistics: A Report of US and Global Data From the American Heart Association. Circulation. 2025;151:e41–e660.

2. Desta L, Jernberg T, Löfman I, Hofman-Bang C, Hagerman I, Spaak J, Persson H. Incidence, temporal trends, and prognostic impact of heart failure complicating acute myocardial infarction. The SWEDEHEART Registry (Swedish Web-System for Enhancement and Development of Evidence-Based Care in Heart Disease Evaluated According to Recommended Therapies): a study of 199,851 patients admitted with index acute myocardial infarctions, 1996 to 2008. JACC Heart Fail. 2015;3:234–242.

3. Heusch G, Libby P, Gersh B, Yellon D, Böhm M, Lopaschuk G, Opie L. Cardiovascular remodelling in coronary artery disease and heart failure. Lancet. 2014;383:1933–1943.

4. Frantz S, Hundertmark MJ, Schulz-Menger J, Bengel FM, Bauersachs J. Left ventricular remodelling post-myocardial infarction: pathophysiology, imaging, and novel therapies. Eur Heart J. 2022;43:2549–2561.

5. Zhang J, Bolli R, Garry DJ, Marbán E, Menasché P, Zimmermann WH, Kamp TJ, Wu JC, Dzau VJ. Basic and Translational Research in Cardiac Repair and Regeneration: JACC State-of-the-Art Review. J Am Coll Cardiol. 2021;78:2092–2105.

6. Menasché P. Cell therapy trials for heart regeneration - lessons learned and future directions. Nat Rev Cardiol. 2018;15:659–671.

7. Petrus-Reurer S, Romano M, Howlett S, Jones JL, Lombardi G, Saeb-Parsy K. Immunological considerations and challenges for regenerative cellular therapies. Commun Biol. 2021;4:798.

8. Chatenoud L. Immunology. Teaching the immune system “self” respect and tolerance. Science. 2014;344:1343–1344.

9. Brown CC, Rudensky AY. Spatiotemporal regulation of peripheral T cell tolerance. Science. 2023;380:472–478.

10. Zakrzewski JL, van den Brink MR, Hubbell JA. Overcoming immunological barriers in regenerative medicine. Nat Biotechnol. 2014;32:786–794.

11. Jiang Y, Sun SJ, Zhen Z, Wei R, Zhang N, Liao SY, Tse HF. Myocardial repair of bioengineered cardiac patches with decellularized placental scaffold and human-induced pluripotent stem cells in a rat model of myocardial infarction. Stem Cell Res Ther. 2021;12:13.

12. Lanza R, Russell DW, Nagy A. Engineering universal cells that evade immune detection. Nat Rev Immunol. 2019;19:723–733

13. Sun SJ, Lai WH, Jiang Y, Zhen Z, Wei R, Lian Q, Liao SY, Tse HF. Immunomodulation by systemic administration of human-induced pluripotent stem cell-derived mesenchymal stromal cells to enhance the therapeutic efficacy of cell-based therapy for treatment of myocardial infarction. Theranostics. 2021;11:1641–1654.

14. Sun SJ, Li F, Dong M, Liang WH, Lai WH, Ho WI, Wei R, Huang Y, Liao SY, Tse HF. Repeated intravenous administration of hiPSC-MSCs enhance the efficacy of cell-based therapy in tissue regeneration. Commun Biol. 2022;5:867.

15. Ferreira LMR, Meissner TB, Tilburgs T, Strominger JL. HLA-G: At the Interface of Maternal-Fetal Tolerance. Trends Immunol. 2017;38:272–286.

16. Nikodemova M and Watters JJ. Outbred ICR/CD1 mice display more severe neuroinflammation mediated by microglial TLR4/CD14 activation than inbred C57BL/6 mice. Neuroscience. 2011;190:67–74.

17. Zacchigna, S., A. Paldino, I. Falcao-Pires, E. P. Daskalopoulos, M. Dal Ferro, S. Vodret, P. Lesizza, A. Cannata, D. Miranda-Silva, A. P. Lourenco, B. Pinamonti, G. Sinagra, F. Weinberger, T. Eschenhagen, L. Carrier, I. Kehat, C. G. Tocchetti, M. Russo, A. Ghigo, J. Cimino, E. Hirsch, D. Dawson, M. Ciccarelli, M. Oliveti, W. A. Linke, I. Cuijpers, S. Heymans, N. Hamdani, M. de Boer, D. J. Duncker, D. Kuster, J. van der Velden, C. Beauloye, L. Bertrand, M. Mayr, M. Giacca, F. Leuschner, J. Backs, and T. Thum. Towards standardization of echocardiography for the evaluation of left ventricular function in adult rodents: a position paper of the ESC Working Group on Myocardial Function. Cardiovascular Research. 2021;117:43–59.

18. Ferreira LM, Meissner TB, Mikkelsen TS, Mallard W, O’Donnell CW, Tilburgs T, Gomes HA, Camahort R, Sherwood RI, Gifford DK, Rinn JL, Cowan CA, Strominger JL. A distant trophoblast-specific enhancer controls HLA-G expression at the maternal-fetal interface. Proc Natl Acad Sci USA. 2016;113:5364–5369.

19. Lintao RCV, Richardson LS, Kammala AK, Chapa J, Yunque-Yap DA, Khanipov K, Golovko G, Dalmacio LMM, Menon R. PGRMC2 and HLA-G regulate immune homeostasis in a microphysiological model of human maternal-fetal membrane interface. Commun Biol. 2024;7:1041.

20. Takashi Yamagami, Chad Sanada, Heinz Wiendl, Esmail D. Zanjani, Christopher D. Porada, Graça Almeida-Porada; Expression of HLA-G1 and G5 Enables Human Mesenchymal Stem Cells To Engraft at High Levels across Xenogeneic Barriers Following Transplantation into Immunocompetent Recipients. Blood. 2007;110:1195.

21. Zoehler B, Fracaro L, Boldrini-Leite LM, da Silva JS, Travers PJ, Brofman PRS, Bicalho MDG, Senegaglia AC. HLA-G and CD152 Expression Levels Encourage the Use of Umbilical Cord Tissue-Derived Mesenchymal Stromal Cells as an Alternative for Immunosuppressive Therapy. Cells. 2022;11:1339.

22. Zakrzewski JL, van den Brink MR, Hubbell JA. Overcoming immunological barriers in regenerative medicine. Nat Biotechnol. 2014;32:786–794.

23. Raeber ME, Sahin D, Karakus U, Boyman O. A systematic review of interleukin-2-based immunotherapies in clinical trials for cancer and autoimmune diseases. EBioMedicine. 2023;90:104539.

24. Song N, Scholtemeijer M, Shah K. Mesenchymal Stem Cell Immunomodulation: Mechanisms and Therapeutic Potential. Trends Pharmacol Sci. 2020;41:653–664.

25. Bartolucci J, Verdugo FJ, González PL, Larrea RE, Abarzua E, Goset C, Rojo P, Palma I, Lamich R, Pedreros PA, Valdivia G, Lopez VM, Nazzal C, Alcayaga-Miranda F, Cuenca J, Brobeck MJ, Patel AN, Figueroa FE, Khoury M. Safety and Efficacy of the Intravenous Infusion of Umbilical Cord Mesenchymal Stem Cells in Patients With Heart Failure: A Phase 1/2 Randomized Controlled Trial (RIMECARD Trial [Randomized Clinical Trial of Intravenous Infusion Umbilical Cord Mesenchymal Stem Cells on Cardiopathy]). Circ Res. 2017;121:1192–1204.

26. Luger D, Lipinski MJ, Westman PC, Glover DK, Dimastromatteo J, Frias JC, Albelda MT, Sikora S, Kharazi A, Vertelov G, Waksman R, Epstein SE. Intravenously Delivered Mesenchymal Stem Cells: Systemic Anti-Inflammatory Effects Improve Left Ventricular Dysfunction in Acute Myocardial Infarction and Ischemic Cardiomyopathy. Circ Res. 2017;120:1598–1613.

27. Cheng YC, Hsieh ML, Lin CJ, Chang CMC, Huang CY, Puntney R, Wu Moy A, Ting CY, Herr Chan DZ, Nicholson MW, Lin PJ, Chen HC, Kim GC, Zhang J, Coonen J, Basu P, Simmons HA, Liu YW, Hacker TA, Kamp TJ, Hsieh PCH. Combined Treatment of Human Induced Pluripotent Stem Cell-Derived Cardiomyocytes and Endothelial Cells Regenerate the Infarcted Heart in Mice and Non-Human Primates. Circulation. 2023;148:1395–1409.

28. Papuchova H, Kshirsagar S, Xu L, Bougleux Gomes HA, Li Q, Iyer V, Norwitz ER, Strominger JL, Tilburgs T. Three types of HLA-G+ extravillous trophoblasts that have distinct immune regulatory properties. Proc Natl Acad Sci USA. 2020;117:15772–15777.

